# Social network dynamics and gut microbiota composition during alpha male challenges in *Colobus vellerosus*

**DOI:** 10.1101/2023.08.22.554322

**Authors:** Shelby Samartino, Diana Christie, Anna Penna, Pascale Sicotte, Nelson Ting, Eva Wikberg

## Abstract

The gut microbiota of group-living animals is strongly influenced by their social interactions, but it is unclear how it responds to social instability. We investigated whether social instability associated with the immigration of new males and challenges to the alpha male position could explain differences in the gut microbiota in adult female *Colobus vellerosus* at Boabeng-Fiema, Ghana. During May-August 2007 and May 2008-May 2009, we collected: 1) 53 fecal samples from adult females in 8 social groups for v4 16S rRNA sequencing to determine gut microbiota composition; and 2) demographic and behavioral data *ad libitum* to determine male immigration, challenges to the alpha male position, and infant births and deaths. We estimated Sørensen and Bray-Curtis beta diversity indices (i.e., between-sample microbiome variation), and they were predicted by year, alpha male stability, group identity, age, and individual identity. We then created 1-m proximity networks using detailed behavioral data via focal follows of 19 adult females in 3 of these groups. Yearly 1-m proximity ties predicted adult female beta-diversity in the two socially stable groups. An alpha male takeover in the third group was associated with infant mortality and temporal variation in proximity networks. Beta-diversity among adult females was predicted by similarity in infant loss status and short-term (rather than yearly) 1-m proximity ties. Although the mechanism driving this association needs to be further investigated in future studies, our findings indicate that alpha male takeovers and social stability are associated with gut microbiota variation and highlight the importance of taking demographic and social network dynamics into account.

## Introduction

The gut microbiome is made up of a combination of bacteria, viruses, archaea, and eukaryotes that can alter host nutritional status, immune function, and behavior (McFall-Ngai et al., 2013). The gut microbiota can have important health and fitness consequences for animal hosts in wild (Koch & Schmid-Hempel, 2011) and captive settings (Turnbaugh et al., 2006), and the gut microbiota composition differs between healthy people and people with certain diseases (Bäckhed et al., 2012). A growing number of studies indicate that social interactions may help maintain a microbiota that promotes host health, survival, and reproduction (Archie & Tung, 2015; Baniel & Charpentier, 2022; Björk et al., 2019; Lombardo, 2008; Sarkar et al., 2020).

Because social interactions occur more often within than between groups, each social group has a unique microbiome in many group-living animal populations, including humans (Lax et al., 2014; Song et al., 2013; Yatsunenko et al., 2012), non-human primates (Amato et al., 2017; Goodfellow et al., 2019; Grieneisen et al., 2017; Orkin, Webb, et al., 2019; Perofsky et al., 2017; Raulo et al., 2017; Tung et al., 2015; Wikberg et al., 2020), carnivores (Leclaire et al., 2014; Theis et al., 2013), ungulates (Antwis et al., 2018), and mice (Raulo et al., 2021). Within social groups, patterns of social interactions also predict gut microbiota similarities among group members (Amato et al., 2017; Antwis et al., 2018; Dill-McFarland et al., 2019; Grieneisen et al., 2017; Moeller et al., 2016; Perofsky et al., 2017; Raulo et al., 2017, 2021; Tung et al., 2015; Wikberg et al., 2020). In some of these populations, the similarity in gut microbiota composition between group members is better explained by social interactions than by similarity in diet or relatedness (Amato et al., 2017; Moeller et al., 2016; Perofsky et al., 2017; Raulo et al., 2017; Tung et al., 2015; Wikberg et al., 2020). Direct contact between social partners like grooming (Tung et al., 2015), and indirect contact like touching shared surfaces (Lax et al., 2014) may transmit microbes between hosts. Via these social interactions and contexts, individual communities of microbes are connected, referred to as “the social microbiome” (Sarkar et al., 2020).

Despite the importance of social interactions to gut microbiome composition, very few studies have investigated the impact of social instability. Indeed, for simplicity or due to the lack of sufficient data on shorter time scales, social networks are often quantified as static during the entire study period (Pinter-Wollman et al., 2014). The relationship between social bonds and the gut microbiota when social networks are altered is thus unclear. Immigration events have been suggested to lead to social transmission of microbes between new social partners, which suggests that changing social ties restructures the gut microbiota (Grieneisen et al., 2017; Perofsky et al., 2021). Notably, these studies show that the more time immigrant male baboons (*Papio cynocephalus*) or immigrant sifakas (*Propithecus verreaux*i) reside within a new social group, the more similar their gut microbiota becomes to the members of that group (Grieneisen et al., 2017; Perofsky et al., 2021).

The addition of a new group member via immigration may alter the group’s social network in more ways than simply introducing a new potential social partner. For example, male immigration events that are associated with takeovers of the alpha male position often lead to targeted attacks on infants in white-faced capuchins (*Cebus imitator*), and infanticide can be a major source of infant mortality (Brasington et al., 2017). Immigration events, elevated infanticide risk, and associated social instability are stressful situations and are associated with elevated glucocorticoid levels in chacma baboons (*Papio hamadryas ursinus*) and rhesus macaques (*Mucaca mulatta)* (Engh et al., 2006a, 2006b; Capitanio and Cole, 2015). Although glucocorticoid levels have an impact on gut microbiota composition (Allen-Blevins et al., 2017; Hickmott et al., 2022; Marin et al., 2017; Petrullo et al., 2022; Stothart et al., 2016; but see Rudolph et al., 2022), the direction of the relationship between glucocorticoids and gut microbiota composition varies between studies. For example, higher glucocorticoid levels were associated with increased gut microbiota alpha diversity (i.e., diversity in the gut microbiota composition within a sample) in a study on bonobos (*Pan paniscus*) (Hickmott et al., 2022), while it was associated with decreased alpha diversity in a study of North American red squirrels (*Tamiasciurus hudsonicus*) (Petrullo et al., 2022).

Our study species, the black-and-white colobus monkey (*Colobus vellerosus*), is one of several rare species of arboreal leaf-eating monkeys distributed across the forested regions of the African tropics (Wikberg et al., 2022) and is closely related to guerezas (*Colobus guereza*) and western black-and-white colobus (*Colobus polykomos*) (Ting, 2008). To break down hard-to-digest items in their primarily folivorous diet (Saj & Sicotte, 2007; Teichroeb & Sicotte, 2009), they rely on behavioral and physiological traits, and their gut microbiota (Amato et al., 2016; Lambert, 1998). In the *Colobus vellerosus* population at Boabeng-Fiema, Ghana, all males disperse from their natal group (Teichroeb et al., 2011) while approximately half of the females disperse, often due to high infanticide risk (Sicotte et al., 2017; Teichroeb et al., 2009; Wikberg et al., 2012). Males immigrate to new social groups alone or with other males, and this event is often followed by prolonged periods of intense male-male aggression and/or attacks on young infants (Saj & Sicotte, 2005; Teichroeb & Sicotte, 2008). Changes in male group membership and alpha male takeovers are well documented, and often associated with infant-targeted attacks by males and almost 40% of infant mortality is due to infanticide in some periods (Saj & Sicotte, 2005; Sicotte et al., 2017; Teichroeb et al., 2012; Teichroeb & Sicotte, 2008). Outside these periods of social upheaval, groups have a clear alpha male. Female dominance hierarchies are relatively stable (Wikberg et al., 2013) and intensive female-female aggression mostly occurs during targeted aggression preceding emigration of maturing females (Teichroeb et al., 2009). Colobus monkeys spend a low percentage of their time engaging in direct social interactions (Teichroeb et al., 2003; Wikberg et al., 2014a) possibly due to constraints imposed by their highly folivorous diet (Saj & Sicotte, 2007). Males rarely engage in affiliation with other males or females (Teichroeb et al., 2014; Wikberg et al., 2012). Female colobus spend on average 3% of their time within 1 meter and 0.1% of their time grooming each other, and the social networks are differentiated in most groups (Wikberg et al., 2014a, 2014b, 2015). Despite spending a low percentage of time engaging in social interactions, recent studies suggest that within-group and between-group social connectedness is a better predictor of gut microbiota composition than dietary similarity or relatedness (Wikberg et al., 2020). The Boabeng-Fiema colobus population thus offers an interesting opportunity to investigate how gut microbiota composition may vary with social upheaval.

The aim of this study was to provide a first examination of the association between immigration events and challenges to the alpha male position, infant loss, social network changes, and gut microbiota composition in the population of *Colobus vellerosus* at Boabeng-Fiema. First, we investigated whether alpha male stability (i.e., period with a well-established alpha male versus period when a new immigrant male is aggressively and repeatedly challenging the alpha male) predicts gut microbiota beta diversity (between-sample variation) using samples from two field seasons collected from eight different social groups. As predictor variables, we also included age as it is associated with gut microbiota composition in other study populations (Aivelo et al., 2016; Pafčo et al., 2019; Rudolph et al., 2022; Wei et al., 2022). As suggested by previous studies, we also evaluated the relationship between study period, group ID, and individual ID and gut microbiota composition (Wikberg et al., 2020). We then examined the correlation between the long-term (12 month) and short-term (3 month) 1-m proximity network matrices to assess whether social networks are stable over time in groups with and without alpha male takeovers. For that, we focused on three study groups with detailed behavioral data and gut microbiota samples from the majority of adult females. We predicted that the long-term and short-term networks will be correlated in two study groups (named “BO group” and “OD group”) with a well-established alpha male but not in the group (named “WW group”) which was experiencing alpha male takeovers. We also evaluated which factors predict gut microbiota composition within these three social groups with different social stability. We expected gut microbiota beta-diversity to be negatively correlated with time spent in 1-meter during a one-year time period preceding and overlapping the sampling period (Amato et al., 2017; Perofsky et al., 2017; Tung et al., 2015; Wikberg et al., 2020). Although previous studies reported a strong correlation between beta-diversity and long-term social bond strength (Amato et al., 2017; Perofsky et al., 2017; Tung et al., 2015; Wikberg et al., 2020), gut microbiota composition is sensitive to changes in the social networks (Perofsky et al., 2017). Therefore, we expected a stronger correlation between beta-diversity and short-term rather than long-term proximity networks in the group with alpha male challenges. We also expected that conditions associated with infant loss during alpha male instability will predict gut microbiota beta diversity in WW group. This may be because time periods before or after infant deaths are stressful situations associated with increased glucocorticoid levels (Sapolsky, 2005; Peel et al., 2005; Engh et al., 2006a, 2006b; Capitanio and Cole, 2015; King et al., 2023) that is in turn associated with gut microbiota composition (Allen-Blevins et al., 2017; Hickmott et al., 2022; Marin et al., 2017; Stothart et al., 2016). In OD group with a recent immigrant adult female, we expected gut microbiota beta diversity to be predicted by the dyad’s co-residency status (long-term or short-term co-residents). We also included the predictor variable age similarity and dyadic relatedness in this analysis as these predict gut microbiota composition in other study populations (Reese et al., 2021; Rudolph et al., 2022; Tavalire et al., 2021; Wei et al., 2022; Yatsunenko et al., 2012; Yuan et al., 2015).

## Materials and Methods

We conducted our study on the black-and-white colobus monkeys (*Colobus vellerosus*) at Boabeng-Fiema, Ghana. Pascale Sicotte started studying this population in 2000. Individual recognition of the study animals was achieved sometime between 2000 and 2008. Our research adheres to ASAB ABS Guidelines for the Use of Animals in Research and the laws of Ghana, and data collection was approved by the Boabeng-Fiema Monkey Sanctuary’s management committee, Ghana Wildlife Division and the University of Calgary’s Animal Care Committee (BI 2006-28, BI 2009-25).

Demographic data were collected between May and August 2007 and from May 2008 to April 2009 from up to eight social groups (Table 1). Each group contained 3 to 8 adult females with ages ranging from 5 years to over 15 years. We classified a female as an adult the year she started reproducing. Each study group contained 1-6 adult males (i.e., fully grown with a known or estimated age of at least seven years).

**Table 1.**
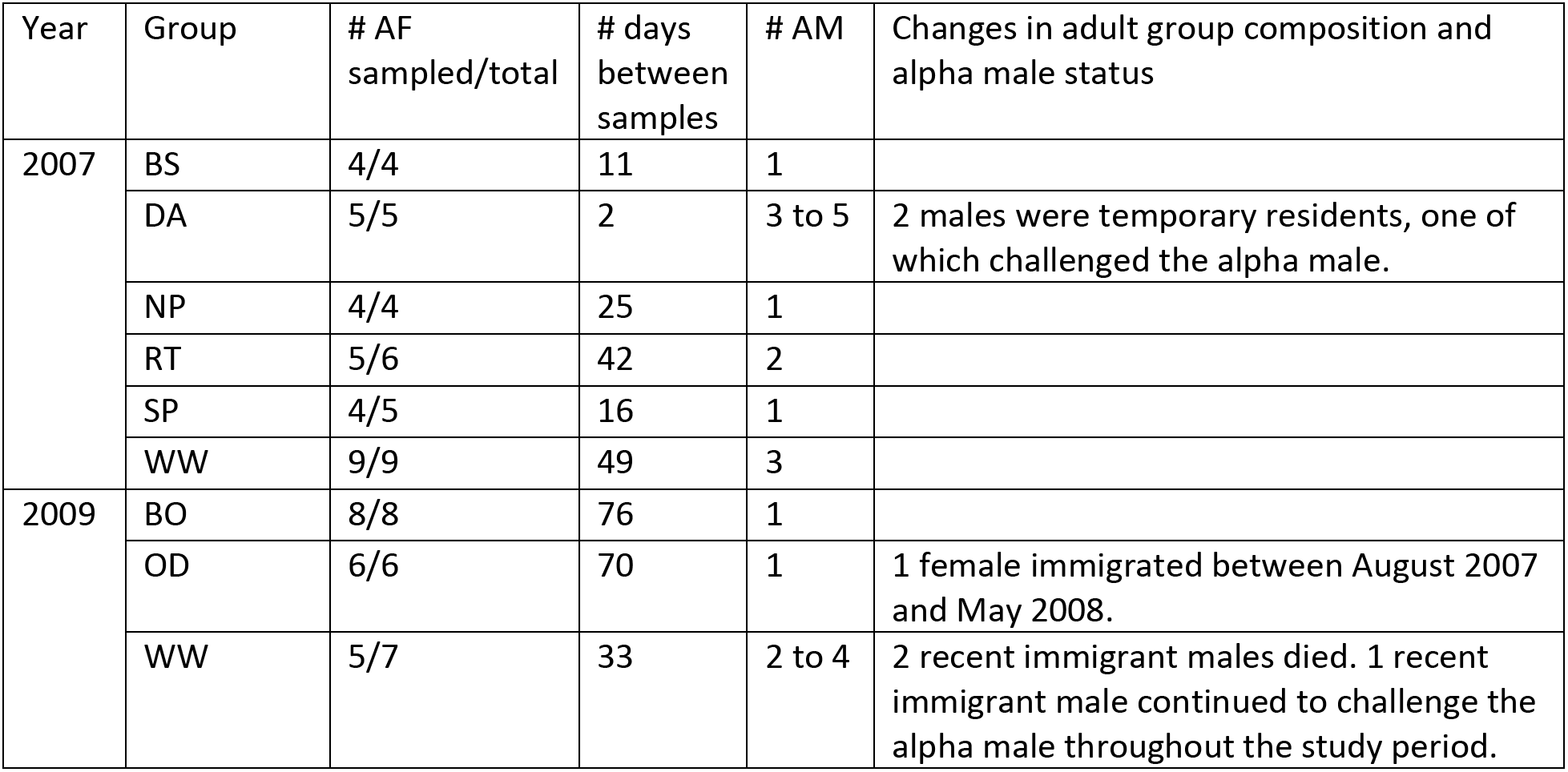
Study period, groups, adult group composition, number of adult females from which we obtained samples for gut microbiome analyses, period during which fecal samples were collected, and changes in male group composition.

Behavioral data were collected from May 2008 to April 2009 during 10-minute focal samples (Altmann, 1974) of adult females and included all social behaviors, social partner identity, and behavior duration. We used point samples collected every 2.5 minutes during the focal follows to record the focal female activity and identities of all individuals within 0, 1, 3, and 5 meters of the focal female. The data collected during focal sampling were used to create adult female interaction matrices with the dyadic percentage of time spent grooming and with the rates of approaches to 1-meter proximity.

We collected fecal samples from the majority of adult females from six groups in the rainy season of 2007 and from three groups in the dry season of 2009 (N = 50 samples) (Table 1). We collected 80% of samples from the rainy season and 70% of samples from the dry season within a 30-day period. We dissolved approximately 1g of feces in 6ml RNAlater. The samples were stored in a fridge at the field site until May 2009 and later transferred to a –20-degree C freezer in the Ting lab. Sample DNA was extracted and genotyped at 17 short tandem repeat loci (STR) as previously described (Wikberg et al., 2012). We calculated dyadic estimated relatedness values (*R*) in MLRelate (Kalinowski et al., 2006) as this method provided the most accurate relatedness estimates in our study population (Wikberg et al., 2012). *R*-values were calculated from STR loci rather than theoretical relatedness (*r*) calculated from pedigrees, because *R*-values predict kinship relatively accurately in our study population (Wikberg et al., 2014b) and they are more accurate than *r* in studies such as ours with limited access to pedigrees (Forstmeier et al., 2012; Robinson et al., 2013). We compared the STR genotypes obtained from two samples collected from the same individual at a different time to ensure samples used in relatedness and gut microbiota analyses were from the correct individual.

For the gut microbial data, we extracted DNA from previously genotyped samples using the QIAamp DNA Stool Mini Kit with a modified protocol (Wikberg et al., 2020). We amplified the V4 hypervariable region of the bacterial 16S ribosomal RNA gene and completed library preparation using the 515F and 806R primers containing 5’ Illumina adapter tails and dual indexing barcodes. The libraries were sequenced as part of a 150bp paired-end sequencing run on the Illumina NextSeq platform. Sequences were demultiplexed, denoised, and assigned as Amplicon Sequence Variants (ASVs) in QIIME2 and DADA2 following current recommendations (Callahan et al., 2017). Taxonomic information was obtained from the SILVA rRNA database (Glöckner et al., 2017). Rare sequences with a frequency less than 0.0005 were filtered out (Wikberg et al., 2020). We also rarified the data to an even read depth of 56,234. We used the vegan R package (Oksanen et al., 2017) to calculate two measures of gut microbiota beta-diversity: Sørensen dissimilarity index, which takes presence and absence of ASVs into account, and Bray-Curtis dissimilarity index, which takes the relative abundance of ASVs into account.

### Data analyses

We first investigated variation in gut microbiota beta diversity using 53 adult female samples collected from 6 social groups during the rainy season in 2007 and 3 social groups during the dry season in 2009. We analyzed the effects of study period (i.e. rainy season 2007 or dry season 2009), alpha male stability, group identity, and individual age on Sørensen dissimilarity index or the Bray-Curtis dissimilarity index of gut microbiota beta diversity by adding these terms sequentially in this order in a permutational multivariate analysis of variance (PERMANOVA). We ran the analysis with 10 000 permutations using the ‘adonis2’ function in the ‘vegan’ R package (Oksanen et al., 2017).

Social bond strength in primates is often measured as the time spent grooming. However, close proximity may be a better predictor of social bond strength in populations with low rates of grooming like in our study population (Wikberg et al., 2014a). To assess whether 1-meter proximity matrix was correlated with the grooming matrix, we used Quadradic Assignment Procedures (QAP) in UCINET (Borgatti et al., 2002). We also analyzed the correlation between the 3-month and the 12-month 1-meter proximity matrices to assess whether social networks were stable. These analyses were conducted with three study groups that we had data on the gut microbiota composition from the dry season in 2009 and up to 12 months of behavioral data from May 2008 to May 2009.

Finally, we further investigated the variation in gut microbiota beta-diversity between adult females in the same social group, again focusing on the three study groups that we had data on the gut microbiota composition from the dry season in 2009 and up to 12 months of behavioral data from May 2008 to May 2009. We created QAP models in UCINET (Borgatti et al., 2002) to examine whether matrices with the Sørensen dissimilarity index or the Bray-Curtis dissimilarity was correlated with the 3-month or the 12-month proximity matrices and with similarity in infant loss status in WW group with alpha male instability. The 3-mo period with interaction data precedes and includes the 17-day fecal sample collection period in WW group. Note that we were unable to do a similar analysis of the correlation between a short-term proximity network and gut microbiota similarity in the two socially stable study groups from which we had behavioral and microbiome data in 2009 because their fecal samples were collected throughout the dry season, up to four months apart. We included similarity in infant loss status as a predictor variable with the mothers who lost their infants shortly before or after the fecal sample collection period being coded as experiencing infant loss while the mother whose infant survived and the adult females that did not have an infant being coded as not experiencing infant loss. We also created QAP models in UCINET (Borgatti et al., 2002) to examine whether matrices with the Sørensen dissimilarity index or the Bray-Curtis dissimilarity index were correlated with the 12-month proximity matrices in BO and OD group. OD group contained one new immigrant adult female, and time spent resident in a social group is associated with gut microbiota composition in other study populations (Grieneisen et al., 2017; Perofsky et al., 2021). Therefore, we also included the dyad’s co-residency status as a predictor variable in OD group, and a dyad was coded as short-term co-residents if the two females had only resided together since May 2008 and long-term co-residents if they had resided together since females were individually identified in May 2006. Because gut microbiota similarity is also predicted by relatedness and age (Reese et al., 2021; Rudolph et al., 2022; Tavalire et al., 2021; Wei et al., 2022; Yatsunenko et al., 2012; Yuan et al., 2015), we also included dyadic *R*-values and age difference as predictor matrices in all models. Age difference was calculated as the absolute number of years that two females differed in age, either using age based on birth records or age estimates done independently by two experienced observers (Wikberg et al., 2014b). Although some group’s proximity networks were structured by kinship in a previous study that included both adult and subadult females (Wikberg et al., 2014a), the current analyses did not contain subadult females, and there was no correlation between close proximity and *R*-values in this set of study females (QAP, p > 0.05). None of the predictor matrices were correlated (QAP, all p > 0.05). We did not correct these p-values for multiple testing following the recommendations for small sample sizes (N = 5-8 females) (Nakagawa, 2004). We consider results to be significant if p < 0.05.

## Results

### Periods with immigration and alpha male challenges

Adult male group composition and alpha male status were stable within each study period and group except from DA group 2007 and WW group 2008-2009. Two new immigrant males were temporarily residing with DA group May to August 2007. The new males were engaged in aggression with each other and with the old resident males. One of the new males appeared to temporarily hold the alpha status, but he disappeared from the group shortly after achieving alpha status. One female gave birth to an infant during this time period, and it disappeared within one month.

In WW group, three adult males had immigrated when observers were absent sometime between August 2007 and May 2008, and two of these males died in July and August 2008. One male died after receiving a large abdominal wound while the other male died when he fell from a tree. The events leading up to their deaths were not observed; however, the new immigrant males were frequently involved in aggression with each other and the previous alpha male. The surviving new immigrant male and the previous alpha male were aggressive towards each other, and the alpha male status remained unclear or frequently changed throughout the 2008-2009 study period. In WW group, four infants were born between November 2008 and February 2009. One infant survived until it reached juvenescence. The other three infants only survived for approximately one month. We did not observe what happened to these infants, but the mothers had new scars after their infants went missing.

Sometime between August 2007 and May 2008 when observers were absent, one adult female immigrated into OD group. She appeared to be fully integrated in the group when the 2008-2009 field season began, and we did not observe her receiving targeted aggression from the other females. In BO group, no females immigrated after it became a study group in 2008, but we lack long-term data on female immigration status. There were no females immigrating to the other groups during the study period 2007-2009.

### Predictors of gut microbiota similarity on a population level

First, we analyzed our full data set that included samples collected from eight groups during the rainy season 2007 and/or the dry season 2009. Based on a principle of components analysis, the first two axes explained 13.1% and 6.9% of the variation using Sørensen index of gut microbiota dissimilarity (Fig. 1). The samples are roughly clustered based on year and season (i.e., dry season 2009 vs. rainy season 2007) and whether or not there was stability in the group’s alpha male position (Fig. 1).

**Figure 1.**
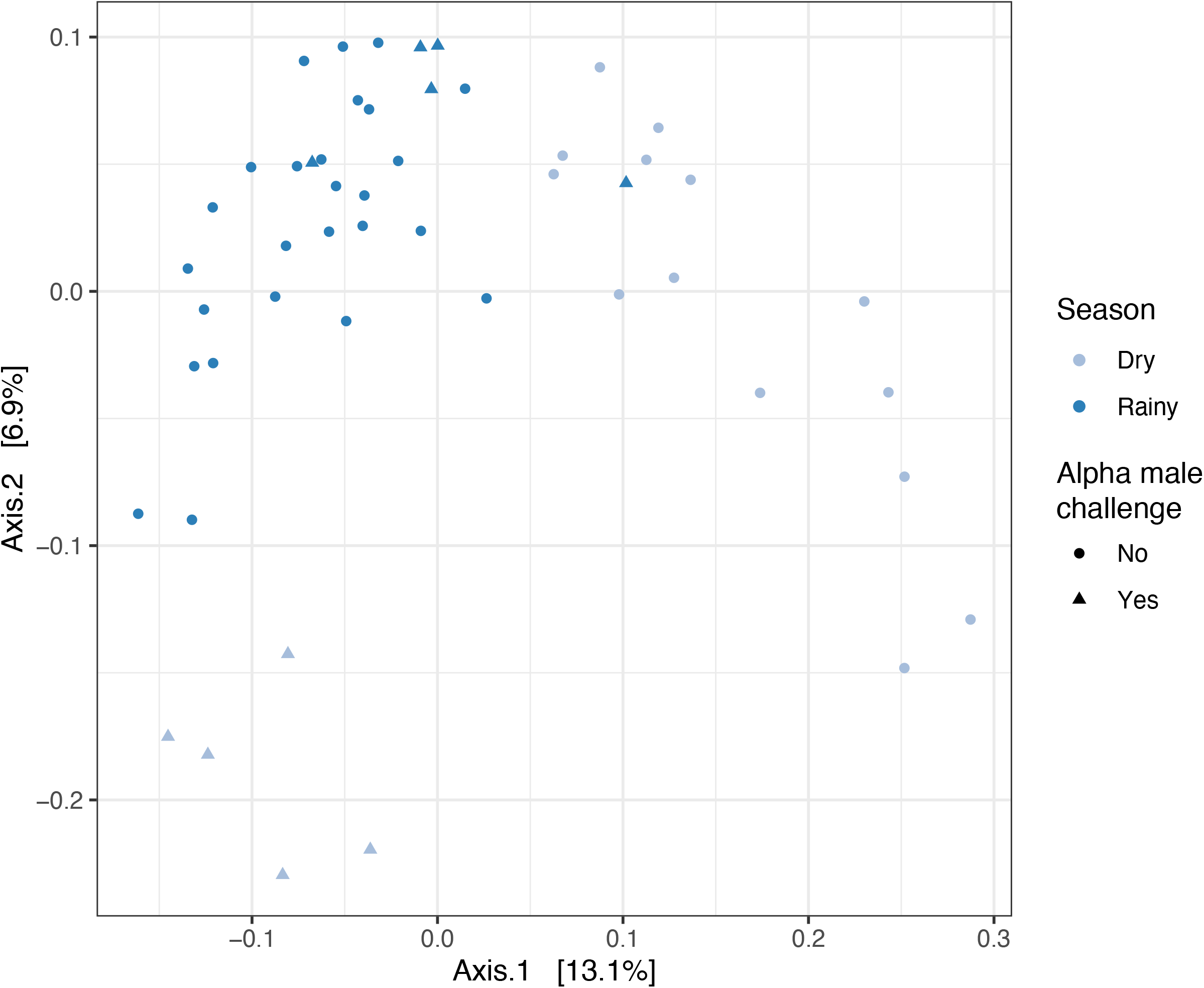
Principal coordinates analysis of Sørensen beta diversity index with shade indicating season (dark blue = rainy season 2007, light blue = dry season 2009) and shape illustrating whether the alpha male position was stable (stable = circles, unstable = triangles).

In this data set, gut microbiota beta diversity was predicted by study period, alpha male stability, group identity, and individual age (PERMANOVA, N = 50 samples, Table 2). Alpha male stability explained 4% of the variation in Sørensen index and Bray-Curtis index of beta diversity. Year and age explained 2-8% percentage of the variation in Sørensen index and Bray-Curtis index of beta diversity. Large percentages of the variation in Sørensen index and Bray-Curtis index of beta diversity were explained by group ID (21-31%) (Table 2).

**Table 2.**
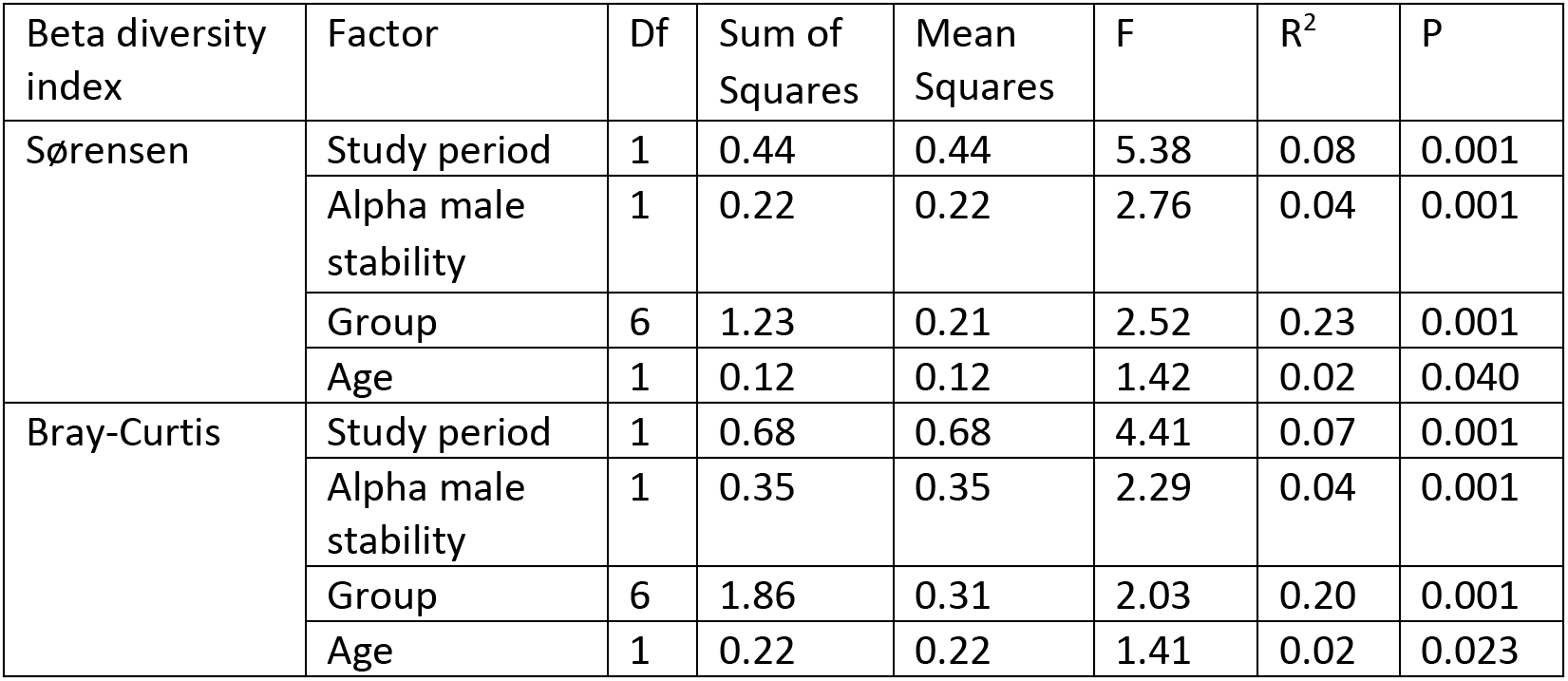
PERMANOVA results for beta diversity index of gut microbiome dissimilarity.

*Female social interaction matrices and predictors of gut microbiota similarity on a group level* Next, we focused on three study groups from which we had collected detailed social interaction data and had sampled the majority of adult females in the dry season 2009. In BO and OD group with no challenges to the alpha male position, the yearly 1-meter proximity matrix was correlated with the yearly grooming matrix (QAP: r_BO_ = 0.43, r_OD_ = 0.67, p < 0.050) and with the dry season proximity matrix (QAP: r_BO_ = 0.75, r_OD_ = 0.65, p < 0.010). This indicates that the 1-meter proximity matrix was a good indicator of grooming bond strength and that social bonds were relatively stable during the study year in BO and OD group. In WW group in which the alpha male was being challenged, the proximity matrix created with data collected during 12 months was not correlated with the proximity matrix for the 3-month period preceding and overlapping with the sample collection period (QAP: r = –0.08, p = 0.375), indicating that the social network in WW group was changing over time. There was a trend towards a strong correlation between the proximity and grooming matrices in this 3-month period even though the grooming matrix was scant (QAP: r = 0.62, p = 0.052). This indicates that the short-term proximity network was a relatively good predictor of short-term grooming bond strength in WW group.

For each group separately, we investigated whether within-group gut microbiota dissimilarity was explained by 1-meter proximity, relatedness, age differences, co-residency status (for OD group only) and/or infant loss (for WW group only). In WW group in which the alpha male was being challenged, the model with the short-term proximity network was significant and explained 74% of the observed variation in the Bray-Curtis index for gut microbiota dissimilarity (Table 3). Short-term proximity and similarity in infant loss were negatively correlated with the Bray-Curtis index for gut microbiota dissimilarity (QAP; N = 5 females; Table 3). This means that the gut microbiota composition was more similar in dyads with the same infant loss status and in dyads that spent a higher proportion of their time in close proximity during the last three months. There was also a trend for Bray-Curtis index to decrease with age similarity (Table 3). The Bray-Curtis index was not correlated with *R*-values (Table 3). The model with the Sørensen index of gut microbiota dissimilarity and short-term proximity network had a p-value of 0.051. The Sørensen index was negatively correlated with short-term proximity and similarity in infant loss status, and there was a tendency for a negative correlation between Sørensen index and age similarity and *R*-values (Table 3). In this group with social instability in which the short-term and long-term proximity network were not correlated, the models of gut microbiota beta diversity using the yearly proximity matrix as a predictor were not significant (Table 3).

**Table 3.**
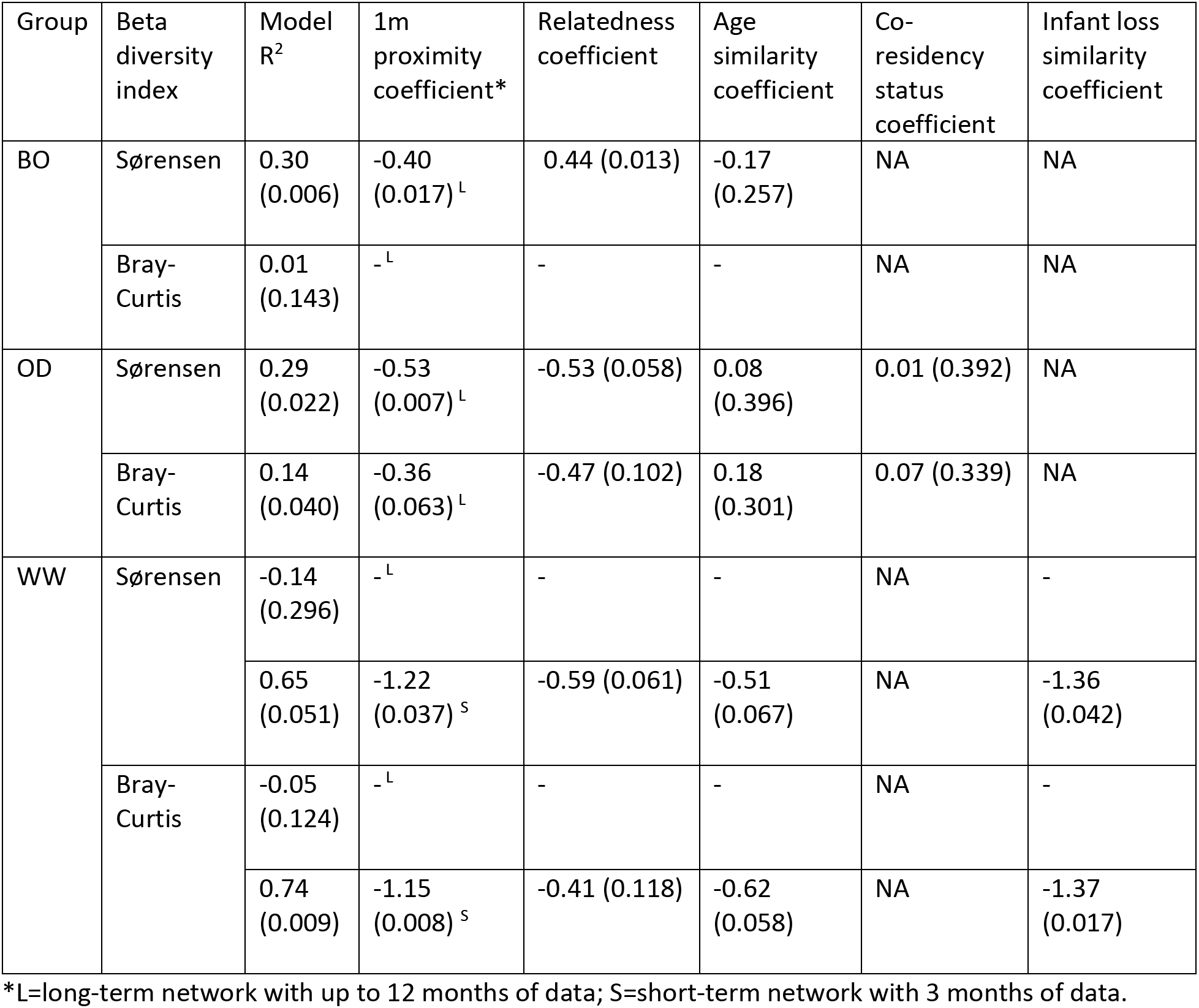
QAP model R^2^ and coefficient estimates with p-values in parentheses for outcome variable Sørensen or Bray-Curtis beta diversity index.

| Group | Beta diversity index | Model R<sup>2</sup> | 1m proximity coefficient* | Relatedness coefficient | Age similarity coefficient | Co-residency status coefficient | Infant loss similarity coefficient |
| --- | --- | --- | --- | --- | --- | --- | --- |
| BO | Sørensen | 0.30 (0.006) | -0.40 (0.017)<sup>L</sup> | 0.44 (0.013) | -0.17 (0.257) | NA | NA |
|  | Bray-Curtis | 0.01 (0.143) | -<sup>L</sup> | - | - | NA | NA |
| OD | Sørensen | 0.29 (0.022) | -0.53 (0.007)<sup>L</sup> | -0.53 (0.058) | 0.08 (0.396) | 0.01 (0.392) | NA |
|  | Bray-Curtis | 0.14 (0.040) | -0.36 (0.063)<sup>L</sup> | -0.47 (0.102) | 0.18 (0.301) | 0.07 (0.339) | NA |
| WW | Sørensen | -0.14 (0.296) | -<sup>L</sup> | - | - | NA | - |
|  |  | 0.65 (0.051) | -1.22 (0.037)<sup>S</sup> | -0.59 (0.061) | -0.51 (0.067) | NA | -1.36 (0.042) |
|  | Bray-Curtis | -0.05 (0.124) | -<sup>L</sup> | - | - | NA | - |
|  |  | 0.74 (0.009) | -1.15 (0.008)<sup>S</sup> | -0.41 (0.118) | -0.62 (0.058) | NA | -1.37 (0.017) |

\*L=long-term network with up to 12 months of data; S=short-term network with 3 months of data.

In BO group and OD group, the model predicting Sørensen index for gut microbiota dissimilarity was significant in both groups. The models explained 29-30% of the observed variation in the Sørensen index of gut microbiota dissimilarity (QAP; N = 8 females in BO group and N = 6 females in OD group; Table 3). The Sørensen index decreased with yearly proximity in both groups (Table 3). The Sørensen index increased with relatedness in BO group, while there was a trend for it to decrease with relatedness in OD group (Table 3). The Sørensen index was not correlated with age difference in these groups nor co-residency status in OD group (Table 3). The model with Bray-Curtis index of gut microbiota dissimilarity was significant for OD group and explained 14% of the observed variation (QAP; N = 6 females; Table 3). The Bray-Curtis index decreased with yearly proximity, but was not correlated with relatedness, age difference, or co-residency status (Table 3). The model with Bray-Curtis index of gut microbiota dissimilarity was not significant for BO group (QAP; N = 8 females; Table 3).

## Discussion

The aim of this study was to provide a first examination of the association between social instability and gut microbiota composition in *Colobus vellerosus* at Boabeng-Fiema. The results indicated that gut microbiota beta diversity (between-sample dissimilarity) is explained by a combination of factors associated with social instability in addition to other, better known factors such as belonging to a specific social group.

Indeed, in our analysis of the full 2007-2009 data set, a large proportion of the variation in gut microbiota composition is explained by group ID (Leclaire et al., 2014; Springer et al., 2017) Greene et al., 2021) with a smaller proportion of the variation being explained by year and season (Maurice et al., 2015; Orkin, Campos, et al., 2019; Springer et al., 2017) and age (Leclaire et al., 2014; Aivelo et al., 2015; Rudolph et al., 2022). When groups were analyzed separately using samples from the dry season of 2009, we found mixed or no support for the predictions that gut microbiota similarity would increase with age similarity, relatedness, and with time spent co-resident (D’Argenio & Salvatore, 2015; Degnan et al., 2012; Grieneisen et al., 2017; Perofsky et al., 2017; Voreades et al., 2014; Yatsunenko et al., 2012). There was no consistent effect of age similarity on gut microbiota similarity across the three study groups, possibly due to small sample sizes and varying female age structure across groups. In a larger data set from this study population, age does predict gut microbiome composition, particularly for immature individuals (Christie et al., 2022).

There was a trend for more closely related females to have more similar gut microbiotas (based on Sørensen beta diversity index) as predicted in OD and WW group. In contrast, more closely related females in BO group had more dissimilar gut microbiotas, but genetic analyses suggest that some of these females may be immigrants (Wikberg et al., 2012). Some of our study groups contain at least some immigrant females (e.g., BO and OD) while other study groups consist of philopatric females (WW) (Wikberg et al., 2012). Recent immigrants have distinct gut microbiome composition compared to long-term resident individuals in other populations (Grieneisen et al., 2017; Perofsky et al., 2021). Although this was not the case in OD group that contained one recent female immigrant, it is possible that this recent immigrant female who entered the group before May 2008 had previously resided in OD group, based on similarities in facial features with a female that disappeared from the group between August 2006 and May 2007. Also, a female assigned as her mother (based on genetic analysis) has been residing in this group since all group members were individually recognized in 2006 (Wikberg et al., 2012). Our findings highlight how factors may structure the gut microbiota differently between social groups, which may be because the groups differ in kinship composition, immigration status, and length of group tenure. Because the gut microbiota is influenced by a variety of factors, it is necessary to collect data during different study periods, groups, and populations to fully understand this complex ecosystem.

Gut microbiota composition was predicted by factors associated with social instability during periods when the alpha male was being challenged for his position. A small percentage of the variation in gut microbiota composition in samples from two study periods and up to eight groups was explained by periods with and without alpha male challenges. Our within-group analysis from females in a socially unstable group revealed that females experiencing or susceptible to infant loss had more similar gut microbiotas to each other than to females that did not experience infant loss or who were not vulnerable to infanticide. Due to the limited data set, we were unable to conclude what might be driving these relationships. However, for a female, living through alpha male takeovers, which leads to vulnerability to infanticide and possibly losing an infant are stressful events (Engh et al., 2006a, 2006b; King et al., 2023). Glucocorticoids are correlated with microbiome composition of wild primates (Hickmott et al., 2022; Vlčková et al., 2018) and captive and wild rodents (Allen-Blevins et al., 2017; Marin et al., 2017; Stothart et al., 2016). It is therefore possible alpha male takeovers, the infanticide threat, and the death of their own infants elevated female glucocorticoid levels, leading to associated changes in their gut microbiota composition. Also, potential variation in microbiota composition with male attacks on infants could be due to changes in bond strength. The former alpha male, the mother, and other females jointly defend infants from male attacks (Saj & Sicotte, 2005; Teichroeb & Sicotte, 2008). However, we have yet to analyze short-term changes in dyadic bonds in this context. To examine whether one of these mechanisms can explain the association between gut microbiota composition, alpha male challenges, and threats to infants, future studies would need a more frequent gut microbiome sampling and a larger sample of females experiencing and not experiencing infant loss during alpha male takeovers and detailed contextual data regarding social interactions at shorter time scales and glucocorticoid levels. Forthcoming research in our study population indicates that social bonds are stronger when young infants (< 3 month old) are present in a social group, which is mirrored by higher gut microbial similarity in social groups with young infants. This strengthened social bonding among adults occurs during a phase when infants are most vulnerable to infanticide, and its plausible that similar social dynamics are at play in the context of alpha male takeover and infant loss as well.

The short-term 1-m proximity networks were correlated with the yearly 1-m networks in the two groups with well-established alpha males, suggesting that social networks were relatively stable over time in these groups. In contrast, the short-term and the long-term proximity networks were not correlated in WW group with ongoing alpha male challenges. In similar studies examining how social structure changes with alpha male takeovers, longer periods of social upheaval and multi-male takeovers observed in *Colobus vellerosus* led to more female dispersal during unstable periods; however, only female members with no offspring dispersed (Sicotte et al., 2017). Some females with young infants would temporarily travel out of range of their group during takeover periods, but would eventually return and fall victim to infanticide (Saj & Sicotte, 2005; Teichroeb & Sicotte, 2008). Challenges to the lead male member in golden snub-nosed monkeys (*Rhinopithecus roxellana)* were associated with more extrapair mating events following the takeover period, and females were consorting more with neighboring bachelor males outside of their social network, possibly as a reproductive counterstrategy (Qi et al., 2020). Thus, alpha male takeovers may lead to social changes within and between social units. In WW group, the 3-month proximity network rather than the 12-month proximity network predicted gut microbiota dissimilarity, which highlights the importance of taking temporal network changes into account during periods of social upheaval rather than treating networks as static over a longer time periods. An implication of this association between network dynamism and the gut microbiota is that continual investment in social bonds over time is likely key for gut microbiota maintenance and stability (Rimbach et al., 2015; Sarkar et al., 2020).

Sarkar and colleagues (2020) coined the term the “social microbiome” to describe the interconnectedness of individual microbial communities within an animal social group. Although the social microbiome may buffer against destabilization of an individual’s microbiota (Sarkar et al., 2020), we have a poor understanding of what happens to the social microbiome when all group members’ microbiomes may be changing under stressful situations like alpha male takeovers that affect the entire social group. Although the composition of the gut microbiota can change rapidly with experimental alteration in diet (David et al., 2013; Greene et al., 2018; Michl et al., 2019), the introduction of an infectious disease or pathogen (Aivelo et al., 2016), or habitat disruption (McCord et al., 2014), the gut microbiome may change at a similar pace with the disruption or restructuring of social networks. It is then possible that the microbiome composition of not only individual group members but of the social microbiome changes. To improve our understanding of the relationships between social network dynamics, individual gut microbiota composition, and the social microbiome, future studies should frequently collect samples before, during, and after periods with social upheaval and analyze these samples with methods allowing for strain tracking and more detailed analyses of social transmission of gut microbes (Björk et al., 2019; Sarkar et al., 2020).

## Acknowledgements

We would like to thank the BFMS management committee, Ghana Wildlife Division, and the University of Calgary’s Life and Environmental Sciences Animal Care Committee for permission to conduct this study; Fernando Campos, Teresa Holmes, Robert Koranteng, and Pepra Samuels for field data collection and data management; Alberta Innovates Technology Futures, American Society of Primatologists, Animal Behaviour Society, International Primatological Society, Leakey Foundation, Natural Sciences and Engineering Research Council of Canada, Sweden-America Foundation, Research Services at the University of Calgary, University of Oregon META Center for Systems Biology, and Wenner-Gren Foundation for funding.

## Data Accessibility Statement

The data will be uploaded to FigShare.

## Author contribution

SS, DC, AP, PS, NT, and ECW contributed to writing the manuscript. SS, DC, ECW analysed the data. DC and ECW conducted the DNA extractions. ECW, PS, and NT designed and funded the study.

## Notes

### Competing Interest Statement

The authors have declared no competing interest.

